# Resolving Biology Beyond the Diffraction Limit with Single-Molecule Localization Microscopy

**DOI:** 10.1101/094839

**Authors:** Nafiseh Rafiei, Daniel Nino, Joshua N. Milstein

## Abstract

Optical imaging provides a window into the microscopic world, but the level of observable detail is ultimately limited by the wavelength of light being employed. By harnessing the physics of photoswitchable dyes and fluorescent proteins, single-molecule localization microscopy (SMLM) provides a window into the nano-world of biology. This mini-review article provides a short overview of SMLM and discusses some of its prospects for the future.

## INTRODUCTION

Optical imaging provides a window into the microscopic world, but the level of observable detail is ultimately limited by the wavelength of light being employed. This “resolution limit” or “diffraction limit” results because light diffracts as it passes through an aperture, such as the objective of a microscope. The minimum separation distance at which two point sources of light are distinguishable can be quantified by the Abbe resolution limit:

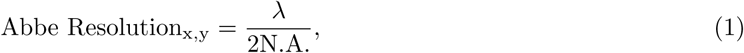

where λ is the wavelength of light and N.A. is the numerical aperture of the imaging lens. ^[1]^ In practical terms, using visible light and a high-N.A. objective, we are limited to resolving structural detail on the order of hundreds of nanometers. This was thought to be a fundamental limit of light microscopy, but in the past decade a number of ways to move beyond the diffraction limit have been devised. It's not that physicists^1^ figured out how to break the diffraction barrier, rather they did what clever scientists do when faced with an insurmountable obstacle. They found a way to get around it.

## SINGLE-MOLECULE IMAGING

A number of techniques have recently been developed to circumvent the diffraction limit. They come bearing a variety of acronyms such as structured illumination microscopy (SIM), ^[2,3]^ stimulated emission depletion (STED) microscopy,^[4]^ photo-activated localization microscopy (PALM),^[5]^ direct stochastic optical reconstruction microscopy (dSTORM),^[6]^ and so on. Here we will focus on techniques like the last two in this list, PALM and dSTORM, which rely upon imaging photoswitchable fluorescent molecules. We refer to these techniques (there are a dozen or so more) as single-molecule localization microscopy so that the reader has to only remember one acronym (SMLM).

SMLM can produce images of structural detail an order-of-magnitude finer than diffraction limited microscopy. The technique relies upon precisely locating the position of single, fluorescent labels (Figure 1). If the system being imaged only contained a single label, say an organic dye, the intensity distribution (or point-spread function (PSF)) of the dye would essentially be an Airy pattern. The central maximum of an Airy pattern is well approximated by a Gaussian, which may be fit to the PSF to obtain the spatial coordinates of the dye. In fact, the precision of this measurement, or the “localization precision”, is primarily limited by the number of photons emitted from the dye, and scales like 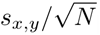, where *s*_*x*,*y*_ is the diffraction limited half-width of the PSF (i.e., the standard deviation of the fitted Gaussian) and *N* is the number of collected photons. A more accurate quantification of the localization precision is given by the following formula:^[7]^

**Figure 1:**
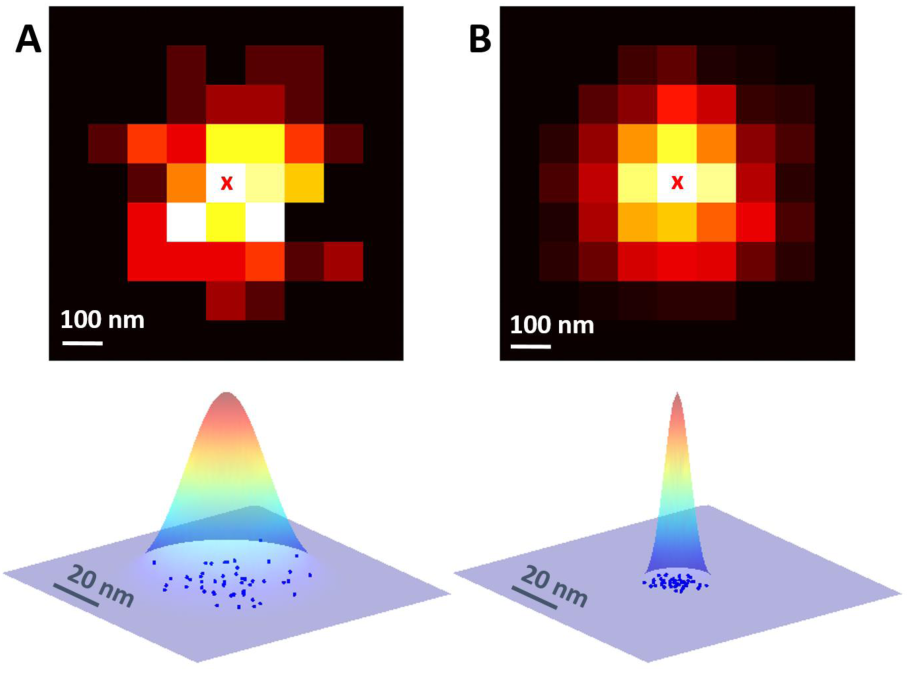
(Top) Pixelated, camera images of a single fluorophore that emits A) *N* = 100 and B) *N* = 1000 photons. The cross (x) represents the fitted center of the intensity distribution (i.e., a localization). (Bottom) Multiple localizations of the fluorophore are represented by the blue circles and are Gaussian distributed. The width of the distribution (or localization precision) scales like 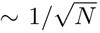. The localization precision is significantly reduced for the brighter fluorophore (A) compared to (B).

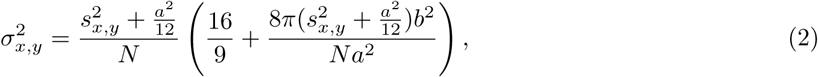

which depends on the pixel size *a* (nm), level of background noise *b* (photons/pixel), standard deviation of the PSF *s*_*x*,*y*_ (nm), and number of collected photons *N*. If there are multiple dyes within close proximity to one another, however, their PSFs will overlap and it will no longer be possible to simply fit the intensity distribution to localize the dyes. The trick, and it really is a trick, is to use photoswitchable dyes so that only a sparse subset of the dyes ever fluoresce at one time (Figure 2). If, on average, only a single fluorophore emits photons at any one time in a diffraction-limited area, then each dye may be localized by fitting the PSF as before. In this context, we often speak of the duty cycle of the dye

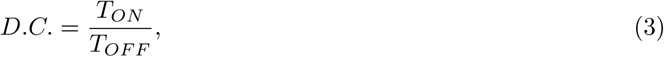

**Figure 2:**
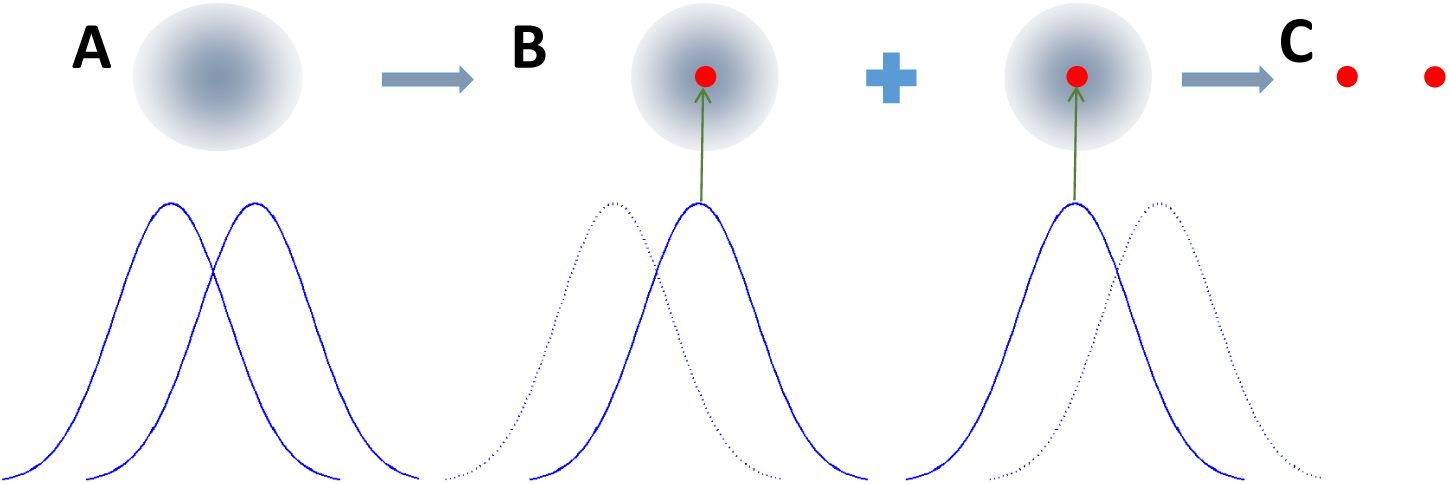
The solid and dotted lines indicate the fluorescent (ON) and non-fluorescent (OFF) states, respectively. A) If the distance between two fluorophores is less than the Abbe limit, they are not resolvable. B) When only one of the fluorophores is in the ON state a 2D Gaussian can be fitted to the PSF to localize the corresponding fluorophore (red circle). C) The process is repeated for the other fluorophore and the positions are assembled to generate a high-resolution image.

which is simply the ratio of the time the dye spends emitting photons (*T_ON_*) to the time it remains dark (*T_OFF_*). On average, so that the dyes’ PSFs don’t overlap, the duty cycle should scale like 1/*M* where *M* is the number of dyes within a diffraction limited area. Many dyes can be tuned to exhibit duty cycles of 10^−4^ − 10^−5^, which is required for high-resolution imaging. ^[8]^

With the current state-of-the-art in SMLM, single dye molecules can be localized with a precision of a few tens of nanometers in the lateral direction. In addition, there are a number of ways to extend SMLM to improve the depth resolution, and thereby perform full 3D imaging,^[9
–11]^ although the localization precision is slightly worse than the lateral case by a factor of 2 or 3 times, dependent on the approach.

## MAKING A FLUOROPHORE BLINK PROPER

As mentioned, the key to SMLM is the ability to actively control the fluorescence emission of photoswitchable fluorophores so that the emission is sufficiently sparse. This can be achieved in a number of ways; for instance, by causing the dyes to intermittently blink through reversibly occupying a long lived dark state, cycling the fluorescence of a portion of the labels between two different wavelengths, or photoactivating a subset then rapidly photobleaching the emitters.

Let's consider PALM imaging,^[5]^ which makes use of inherently photoswitchable fluorescent proteins. A common fluorophore used in PALM is mEOS, which is a green fluorescent protein, but when exposed to near UV light (e.g., 405 nm) a fraction of the fluorophores will behave like a much redder dye and can be excited with a 561 nm laser line. A PALM experiment consists of activating a random subset of the fluorophores into the red channel, imaging those fluorescent proteins, then quickly photobleaching them. A new subset of fluorophores is activated, imaged, bleached, and the cycle repeats. PALM can be performed in fixed or live cells, albeit the requirement of cycling through repeated rounds of localization limits its utility in actively growing, functioning cells. Still, PALM is often used as a way to track proteins within live cells or to obtain rough images of structures that show slow dynamics.

Another approach is to “inactivate” all but a small subset of fluorophores while imaging. Fortunately, most all fluorophores display fluorescence intermittency (i.e., blink) by occasionally transitioning to a triplet or dark state via intersystem crossings before transitioning back to the singlet ground state, often through a non-radiative decay.^[12]^ The time scale of these blinking events, however, is usually on the order of milliseconds or less. The idea of extending the time scale of the fluorescence intermittency is one of the key advances that paved the way for SMLM techniques. As an example, dSTORM, which employs organic dyes such as Alexa-647 or Cy5,^[6]^ makes use of nonfluorescent, long-lived radical ion states beyond the usual triplet state. These dark states, which appear in many commercially available dyes when exposed to millimolar concentrations of thiolating compounds (e.g., beta-mercaptoethanol (BME) or cysteamine (MEA)), can display off times (i.e., when the dye does not fluoresce) of several seconds (Figure 3).

**Figure 3:**
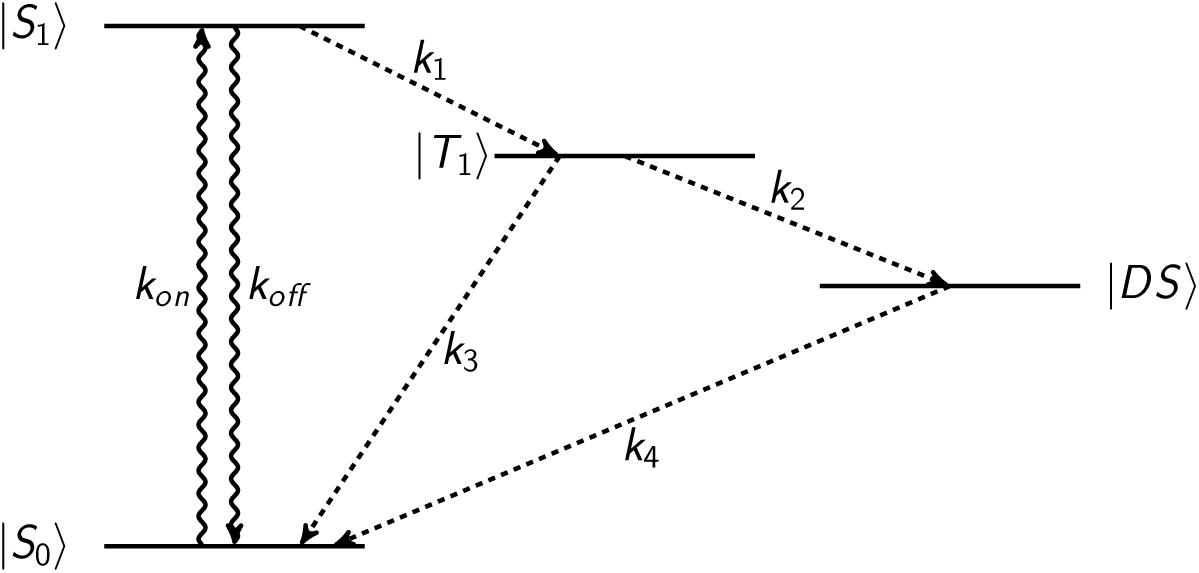
Fluorescence occurs for transitions between the ground |*S*_0_〉 and excited |*S*_1_〉 singlet states. Intersystem crossings can lead to a spin flip transition into the |*T*_1_〉 triplet state, but these states are usually short lived. Long lived, nonradiative, dark states |*DS*〉, however, can be induced in many fluorescent molecules.

## IMAGING THE BACTERIAL PROTEOME

SMLM microscopy is able to provide unprecedented structural detail with visible light microscopy. While the technique has found a range of applications, bacteria are a particularly suitable target because structure within a bacteria was previously inaccessible to light microscopy due to the micron size of these cells. Our lab uses SMLM to try to understand how the organization and packaging of the nucleoid (i.e., bacterial chromosome) affects cell function. In particular, we have focused on the arrangement of highly abundant nucleoid associated proteins (like H-NS, HU, and StpA), which package the bacterial chromosome, similar to histones in eukaryotes, while coordinating the expression of a multiplicity of genes^[15]^

These proteins may serve as environmental sensors that reorganize the chromosome under different environmental conditions, activating networks of genes to assist the cell in adapting to and colonizing its environs. SMLM provides a window into the global organization of these proteins, as well as allowing us to explore stochastic effects such as cell-to-cell variability within a population of bacteria (Figure 4). The technique can also be combined with methods for visualizing chromosomal loci, such as DNA fluorescence in-situ hybridization (FISH), ^[18]^ which allows us to correlate the arrangement of these nucleoid binding proteins with the position of genes within the chromosome.

**Figure 4:**
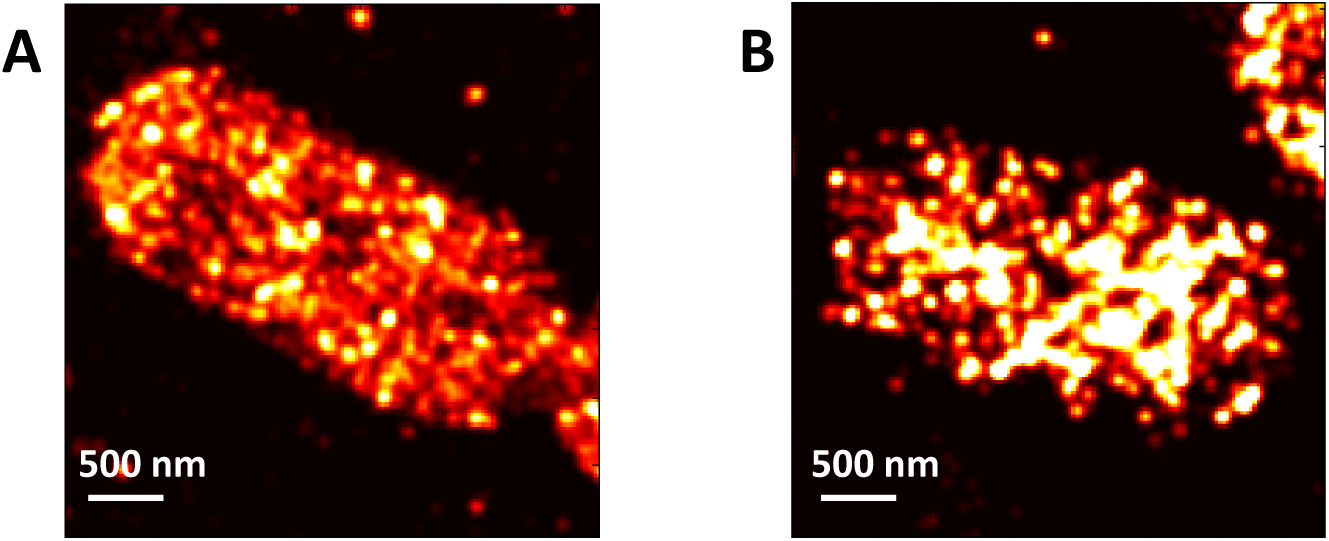
A) PALM image reconstruction of fluorescently labeled H-NS in fixed E. coli bacteria. B) Following cold shock, H-NS is seen to reorganize along the bacterial chromosome into a much more clustered pattern of binding. The localization precision in these images is ~ 30 nm.

## QUANTITATIVE MICROSCOPY AND MOLECULAR COUNTING

Beyond imaging, a promising application of SMLM is as a method to quantify protein or nucleic acid abundance at a single cell level. Molecular counting with SMLM would be particularly powerful in single-cell genomic and proteomic applications. For instance, SMLM should be able to quantify low numbers of amplicons, which would enable a reduction or even elimination of the amplification stage required by current techniques for measuring single-cell DNA or RNA abundance, greatly increasing both the accuracy and reliability of these techniques. Several groups are now working on extracting accurate molecular counts from SMLM data in the hopes that the technique becomes the future gold standard for counting molecules. ^[19
–21]^ Unfortunately, a number of complications still need to be overcome before SMLM is a viable approach to molecular counting. For instance, the photophysics of a fluorophore may be altered by environmental conditions (such as pH) within a cell or by fixation protocols. Moreover, many commonly used SMLM fluorophores are only around 50-60% active,^[20]^ which means that a large portion of the sample will go undetected. This, however, should improve as improved photoswitchable fluorophores are developed. And then there's the issue of generating an accurate table of single-molecule localizations. In practice, especially in samples where there are many aggregates or clusters of proteins, the PSF of the labels will begin to overlap degrading the reliability of the SMLM data set. In some cases, the duty cycle may be further lowered to reduce this overlap, but at the cost of an even even longer acquisition time. Improved localization algorithms ^[7,22]^ as well as increasingly complex models ^[23,24]^ of the fluorophore photophysics may be a better approach toward alleviating this issue.

## CONCLUSION

SMLM has pushed visible light microscopy far beyond the diffraction limit, shedding light on an ever increasing number of biological questions. New and improved fluorophores are continually being developed that are more stable and yield more photons, ever increasing the resolution and speed of this technique. Algorithms for improving the reliability of localization tables (i.e., the acquired list of single-molecule localizations), especially within dense, inhomogeneous samples, are continually being developed as are image analysis tools for extracting quantitative knowledge from SMLM images. Moreover, SMLM can be combined with light-sheet or two-photon imaging to provide super-resolved images deep within samples such as biofilms or the cell nucleus. Finally, quantitative approaches such as molecular counting are gaining traction as an alternative modality for this single-molecule technique, making it increasingly relevant for the rapidly expanding fields of single-cell genomics or proteomics.

## ACKNOWLEDGEMENTS

We thank Dr. Amir Mazouchi for his feedback on this manuscript. Support was provided by the Natural Sciences and Engineering Research Council of Canada and an Early Researcher Award from the Ministry of Research and Innovation.

## Footnotes

1 The 2014 Nobel Prize in Chemistry went to Eric Betzig, Stefan W. Hell and William E. Moerner for developing super resolved microscopy. All three have physics degrees.

